# Computer-Assisted Mitotic Count Using a Deep Learning-based Algorithm Improves Inter-Observer Reproducibility and Accuracy in canine cutaneous mast cell tumors

**DOI:** 10.1101/2021.06.04.446287

**Authors:** Christof A. Bertram, Marc Aubreville, Taryn A. Donovan, Alexander Bartel, Frauke Wilm, Christian Marzahl, Charles-Antoine Assenmacher, Kathrin Becker, Mark Bennett, Sarah Corner, Brieuc Cossic, Daniela Denk, Martina Dettwiler, Beatriz Garcia Gonzalez, Corinne Gurtner, Ann-Kathrin Haverkamp, Annabelle Heier, Annika Lehmbecker, Sophie Merz, Erica L. Noland, Stephanie Plog, Anja Schmidt, Franziska Sebastian, Dodd G. Sledge, Rebecca C. Smedley, Marco Tecilla, Tuddow Thaiwong, Andrea Fuchs-Baumgartinger, Don J. Meuten, Katharina Breininger, Matti Kiupel, Andreas Maier, Robert Klopfleisch

## Abstract

The mitotic count (MC) is an important histological parameter for prognostication of malignant neoplasms. However, it has inter- and intra-observer discrepancies due to difficulties in selecting the region of interest (MC-ROI) and in identifying/classifying mitotic figures (MFs). Recent progress in the field of artificial intelligence has allowed the development of high-performance algorithms that may improve standardization of the MC. As algorithmic predictions are not flawless, the computer-assisted review by pathologists may ensure reliability. In the present study we have compared partial (MC-ROI preselection) and full (additional visualization of MF candidate proposal and display of algorithmic confidence values) computer-assisted MC analysis to the routine (unaided) MC analysis by 23 pathologists for whole slide images of 50 canine cutaneous mast cell tumors (ccMCTs). Algorithmic predictions aimed to assist pathologists in detecting mitotic hotspot locations, reducing omission of MF and improving classification against imposters. The inter-observer consistency for the MC significantly increased with computer assistance (interobserver correlation coefficient, ICC = 0.92) compared to the unaided approach (ICC = 0.70). Classification into prognostic stratifications had a higher accuracy with computer assistance. The algorithmically preselected MC-ROIs had a consistently higher MCs than the manually selected MC-ROIs. Compared to a ground truth (developed with immunohistochemistry for phosphohistone H3), pathologist performance in detecting individual MF was augmented when using computer assistance (F1-score of 0.68 increased to 0.79) with a reduction in false negatives by 38%. The results of this study prove that computer assistance may lead to a more reproducible and accurate MCs in ccMCTs.

---

Proliferation parameters of neoplastic cells correlate with patient prognosis for many tumor types including canine cutaneous mast cell tumors (ccMCTs).^6,9,22,34,38^ The mitotic count (MC) is the only proliferation marker that can be determined quickly and efficiently in standard histological sections stained with hematoxylin and eosin stain (HE).^34^ For ccMCT the MC has been used either as a solitary parameter ^9,32,37^ or as part of tumor grading systems.^25^ If used as a solitary prognostic parameter, a two-tiered system with a MC of 0-5 and a MC > 5 has been evaluated to yield a sensitivity of 32%, 39%, 50% and 55% (high percentage of false negatives) and specificity of 91%, 96%, 98% and 99% (low percentage of false positives) regarding ccMCT-related deaths.^8,9,37,38^ In order to increase sensitivity, other research groups have proposed to use a cut-off value of MC ≥ 2 (sensitivity: 76 and 84%; specificity: 56 and 80% for ccMCT-related death) ^22,37^ or to use a stratification of the MC into three groups (0, 1-7, >7 and 0-1, 2-7, >7, respectively; sensitivity and specificity not available).^18,37^ This data proves that the MC is relevant for prognostication of ccMCT, however, also reveals some variability in the derived results. The question arises whether this variability could be reduced with standardization of the MC methods between different studies.

The MC is mostly defined as the number of mitotic figures (MFs, cells undergoing mitosis visible with microscopy ^17^) within “10 high power field” (equivalent to an area of 2.37 mm²) with the highest mitotic density.^28,29^ Looking at this definition, two potential source of variability for performing the MC become apparent: 1) variable selection of the tumor area and 2) variable detection of MFs. Some studies have proven that it is feasible to perform the MC using digital microscopy and whole slide images (WSIs).^1,7,13,35,42^ As opposed to light microscopy, the concept of enumerating “10 high power fields” (round fields using a microscope at 400x magnification) is extraneous as the area can be accurately measured and labeled in the WSI (with a rectangular field of view). For this study we use the term “mitotic count region of interest” (MC-ROI) for a single, rectangular area with the size of 2.37 mm² in the tumor location.^5,11,28^

Regarding the selection of the MC-ROI, it is common practice to attempt to find a single tumor area with the highest mitotic activity, i.e. a ‘hotspot’.^8,25,28,29,32^ It has often been assumed (but rarely proven, e.g. in human breast cancer ^23^) that the most mitotically active tumor area correlates best with biological behavior of the neoplasm. Of note, it has been shown that the mitotic density varies significantly between different tumor locations in histological sections of ccMCTs and canine mammary carcinomas ^3,5,11^ and that pathologists have some difficulties finding hotspots.^3,11,42^

The pathologist’s capability to identify/classify MFs has been evaluated recently.^15,40,41,43^ Those studies compared the digital MCs of different pathologists in the same MC-ROIs and revealed an overall difference in number of annotated MFs by a factor of 1.5x to 3.3x.^41,43^ This can be attributed to erroneous MF detections related to failure in identification of MF candidates as well as to inaccurate/inconsistent classification of MFs against look-alikes (such as apoptotic bodies, hyperchromatic or deformed nuclei, and inflammatory cells).^17^ Tabata et al. ^35^ have shown that pathologists have a lower accuracy of MF detection when using WSIs (69-74%) as compared to traditional light microscopy (0.8%).

Despite the potential limitations of digital microscopy in detecting MFs, WSIs enable innovative computerized image analysis approaches that have the potential to improve MC reproducibility and accuracy and thereby may improve standardization of the MC methods.^10,14^ Development of high performance image analysis algorithms for MFs has only been possible in the last decade due to the availability of ground-breaking machine learning solutions (especially supervised deep learning) and availability of large-scale datasets.^3,12,39,41^ Regardless of these promising advancements, one of the main points of criticism of deep learning is its ‘black box’ character (i.e. the unavailability of decision criteria) that may lead to failure of identifying algorithmic errors.^10,27^ In order to ensure high reliability, approaches that allow review of the algorithmic predictions by trained pathologists (computer-assisted diagnosis/prognosis) through visualization of algorithmic results as an overlay on the WSI have been recommended for future application.^10^ First computer-assisted MC tools have been recently validated for human pathologists.^7,30^ However, studies that validate the utility of computer assistance for each critical step of performing the MC (see above) and for veterinary tumor histopathology have not been published to date.

The aim of the present study was to compare partially (MC-ROI preselection) and fully (additional MF candidate proposal) computer-assisted MC analysis with routine (unaided) MC analysis in WSIs of ccMCT. We evaluated the assistive value and limitations of computer assistance on the ability of 23 pathologists to perform the overall MC, to determine a count below or above a prognostic cutoff, to select a hotspot MC-ROI, and to identify/classify individual MFs.

## Material and Methods

### Course of studies

For this study, anatomic pathologists (study participants) performed MCs in WSIs of 50 ccMCTs at three stages with different degrees of computer assistance (none, partial and full) (Fig. 1). In stage 1, there was no computer assistance and participants were tasked with screening the WSI manually for MC-ROIs (mitotic hotspots) and annotating all MFs (including atypical MFs) within this area using their “routine” approaches. For stage 2 (partial computer assistance) and stage 3 (strong computer assistance), WSIs were analyzed with a deep learning-based algorithm that detected MFs in the entire tissue section. Based on the algorithmic MF detections, the MC was calculated for each possible tumor location resulting in the MC distribution. The ROI with the highest MC was preselected automatically and presented to the participants in stage 2. For stage 3 (full computer assistance), in addition to the same algorithmic MC-ROI preselection, visualization of the individual MFs and MF look-alike detections (to assist MF *identification*) were provided as an overlay on the WSI along with their corresponding algorithmic confidence values (to assist MF *classification*). Additionally, algorithmic predictions (without review by pathologists) were used for evaluation.

**Figure 1.**
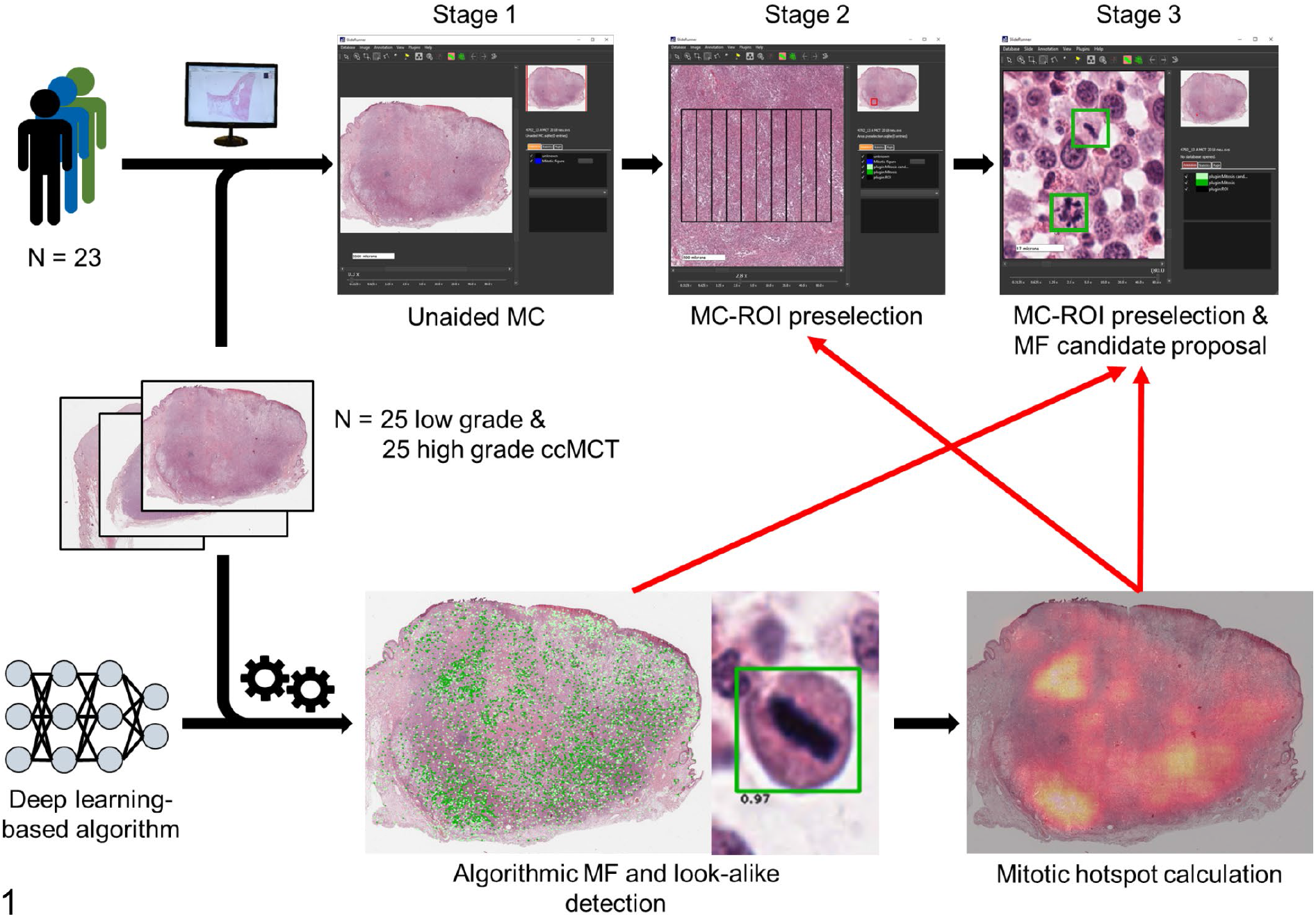
Overview of the course of the study (stage 1-3) with different degrees of computer assistance (red arrows) of the study participants in three examination time points (stages).

Participants were instructed to follow the course of the three stages strictly and to wait at least three days until the next stage. A recall bias was considered negligible due to the increasing amount of computerized information and the very high number of individual MFs. While performing the MCs, participants labeled the enumerated MFs (including atypical MFs) in the exact location of the digital image with a specialized annotation software. This method allows determination of the ability of participants and the ability of the deep learning-based model to identify individual MFs (on the object-level) compared to a gold standard-derived (pHH3 IHC-assisted) ground truth dataset. After the study, participants were asked to fill out an opinion survey.

### Study cases

Random ccMCT cases with an equal distribution of low and high histologic grade ^25^ (based on the original pathology reports) were selected from the archive of the Institute of Veterinary Pathology, Freie Universität Berlin. One tissue block from each of these cases with the largest tumor area was selected. Histological sections were produced from each of the blocks and stained with HE. Glass slides were digitized with a linear scanner (ScanScope CS2; Leica) using default settings. WSIs with one focal scan plane were produced at a magnification of 400x (image resolution: 0.25 μm per pixel). Specimens with an overall very poor tissue preservation (i.e. significant loss of nuclear details in most tumor areas) and cases with a tumor section area of <12 mm² (measured by polygon annotation) were excluded. This procedure was followed until 35 low and 35 high mitotic density were selected. Clinical follow-up (patient outcome) was not considered for this study as the main goal was validation of different MC methods and not to determine prognosis. Local authorities (State Office of Health and Social Affairs of Berlin) approved the use of the samples (approval ID: StN 011/20) for research.

The 70 cases were analyzed with a deep learning-based MF algorithm (see below). Of these, 7 cases (all low grade according to the original reports) had a computerized MC-ROI preselection outside of the tumor area (due to MF predictions mainly in the epidermis, hair follicles, reserve cells of the sebaceous glands or areas of crush artefacts) and were excluded from the study. The remaining 63 cases contained algorithmic MC-ROI preselection within the tumor area and were not further evaluated for exclusion purposes. We randomly excluded 3 additional low grade and 10 high grade cases (according to the original reports) in order to reduce the study set to 25 low grade and 25 high grade ccMCTs. Cases of the study set were randomized in order and labeled with consecutive numbers from 1 to 50. Additionally, a test slide (high grade ccMCT) was provided to allow familiarization of study participants with the annotation software, the study tasks, and the properties of the digital images.

### Study instructions to participants

Twenty-six pathologists from 13 different laboratories volunteered to participate in this study. The study material (WSIs, annotation software, files with algorithmic predictions) was provided, the goal and course of the study was explained and the annotation software was demonstrated (using the test slide) to each participant. For stage 1 participants were instructed to reach for mitotic hotspot MC-ROIs. However, no specific recommendations were given on how to find the “correct” MC-ROI. For all three stages participants were instructed to annotate all MFs in the MC-ROIs with high diligence using their “routine” decision criteria. No specific diagnostic criteria for “correct” identification/classification of MFs (including atypical MFs) were provided in order to validate a realistic diagnostic setting.

### Annotation software and database creation

The open source software SlideRunner ^2^ was used in this study for annotating each enumerated MFs in the precise pixel position in the digital image (object-level). Beyond the ability to view and navigate WSI, this software includes tools for fast image annotations, which are automatically stored in a database. For this study, each participant created their own databases (one for each stage) and annotated each MFs present in the MC-ROIs using a single click tool (saving the x and y coordinate in the WSI). Participants were instructed to pay close attention in placing the annotation in the center of the MFs.

SlideRunner allows integration of software plugins for computerized image analysis and visualization of algorithmic predictions. During specimen selection, all WSIs were analyzed by an MF algorithm (see below) plugin, and predictions of the study cases were saved as separate files that were provided to the participants for visualization in stages 2 and 3.

For each study stage a different SlideRunner package was created, each including a different plugin that enabled visualization of the information relevant for the respective stage as an overlay on the WSI. As intended by the study design, the stage 1 package of the software did not allow visualization of any algorithmic detections. However, this package included a plugin for a rectangular box with the size of exactly 2.37 mm² (aspect ratio of 4:3; width of 1777.6 μm and height of 1333.2 μm). The box was divided into a grid with 9 vertical lines (distance between lines was less than the width of the field of view at 400x magnification), which intended to improve navigation in a meandering pattern. Participants were able to move the box to the desired MC-ROI location by re-centering the box coordinates to the present field of view at any viewing magnification. Due to a software failure, the exact image location of the selected MC-ROIs was not saved in the database. Therefore, we retrospectively determined the approximate MC-ROIs in which the highest number of annotations in the image could be placed. This ensured that the shift between the approximate and the actual selected MC-ROIs were minimal and negligible for our analysis (see below). For 63 cases that did not have an MF annotation (MC = 0), an MC-ROI was not calculated for the respective participant.

The plug-ins of the SlideRunner packages of the two subsequent stages visualized the predictions of the deep learning-based algorithm (Fig. 1). Stage 2 included a plugin with a 2.37 mm²-sized rectangular box (as described above) placed in the region that had the highest density of algorithmic MF detections (partial computer assistance). Unlike stage 1, participants were unable to move the box to another tumor location. The software package for stage 3 displayed the fixed 2.37 mm² box in the same tumor area and additionally visualized all predicted MFs (as dark green boxes) and predicted MF look-alikes (as light green boxes) along with their algorithmic classification scores (“confidence value”). We decided to visualize the MF look-alikes in order to account for potential false negative algorithmic predictions.

After participants finished stage 3, they submitted the three databases to the principal investigators, who verified that all cases had been examined. All annotations that had a center coordinate outside of the 2.37 mm² boxes were deleted as those represented erroneous or unintentional annotations. The participants were anonymized by assigning a random identification number to each participant.

### Image analysis algorithm

Detection results of computerized image analysis were used for stages 2 and 3. Image analysis comprised of two analysis tasks as previously described by Aubreville et al:^5^ 1) a deep learning-based algorithm that detects MFs and 2) the computerized MC density (heat map).

Briefly, the detection of MFs consists of a concatenation of two convolutional neural networks (RetinaNet and ResNet18 architecture).^5^ The first convolutional neural network (object detector) was used to screen the entire WSI for potential MF candidates with high sensitivity and a high processing speed. The second convolutional neural network (patch classifier) was developed to classify small image patches of the detected candidates (by the first neuronal network) into MFs (classification threshold ≥0.5) and MF look-alikes (classification threshold <0.5) with high specificity. The model classification scores (“confidence value”) were extracted from the patch classifier (in order to display along with the algorithmic predictions) and ranged between 0.01 (very unlikely to be a MF) and 1.0 (very likely to be a MF). The models were trained and technically evaluated with an open access dataset with 44,880 MF annotations in 32 ccMCT WSIs that were produced by the same institute that provided the cases for the present study.^12^

The MC density map was derived from calculating the MC, i.e. the number of algorithmic MF detections within a 2.37 mm² box (see above), for every possible center coordinate of a 2.37 mm² box that contains more than 95% tissue.^5,11^ The MC-ROIs for stages 2 and 3 were selected as the image location with the highest density in the MC map.

### pHH3-assisted ground truth

In order to evaluate the pathologists and algorithmic performance on the object level, a ground truth dataset was developed with the assistance of immunohistochemical labeling for phosphohistone H3 (pHH3). pHH3 is a DNA-binding protein that is mostly specific for the mitotic phase of the cell cycle. Histone H3 is phosphorylated in early prophase (still indistinct in HE sections) but already dephosphorylated in telophase.^21^ Based on Tellez et al. ^36^ we established a protocol to de-stain the initial HE-stained sections and immunohistochemically label the same tissue section in order to ensure that the exact same cellular objects were represented in both WSIs. The coverslips of HE-stained sections were removed by incubation in xylene. Subsequent to incubation in a descending alcohol series (99%, 80%) slides were destained under visual control in a 70% alcoholic solution with 0.37% hydrogen chloride. After destaining immunohistochemistry was performed including blockage of endogenous peroxidase (with 10% H_2_O_2_), antigen retrieval by microwave heating (with citrate buffer) and antigens blocked (with goat serum). For the primary antibody we used pHH3 clone E173 (rabbit monoclonal) from Abcam (ab32107, Cambridge, MA, USA) as this product was already used for dogs in a previous study.^31^ Because the HE-stained sections of the study cases were produced at least one year earlier, a somewhat higher antibody concentration of 1:650 (as opposed to new tissue sections with 1:1500) was necessary (established on the excluded cases from this study; see above). The secondary antibody was a goat anti-mouse IgG (H+L) conjugated with alkaline phosphatase and incubated at a dilution of 1:200. 3,3′-Diaminobenzidin (DAB) was used as a chromogen and hematoxylin as counterstain. ccMCTs were used as positive and negative controls. Immunolabeled glass slides were digitized as described above. In 10 of the 50 tissue sections included in the study relevant tissue parts were lost during immunohistochemical processing (if not mounted on adhesive glass slides) or immunohistochemical labeling was nonspecific (internal control compared to the HE staining). Therefore, those cases were excluded from the pHH3-assisted labeling portion of this study.

Following this procedure, we produced two WSIs (one stained with HE and the other labeled with pHH3) from the same tissue section for 40 of the 50 cases. In contrast to using recut tissue sections, this process ensured visualization of the same cellular objects in both WSIs. Automated image registration (according to Jiang et al. ^24^) was performed in order to align the two images so that the tissues matched almost perfectly on a cellular level. Using a newly developed SlideRunner plug-in, it was possible to instantly switch between images with the two staining/labeling methods and therefore easily compare the information for each cell (Fig. 2). The principle investigator (pathologist not involved as a study participant) developed a ground truth dataset for the algorithmically selected MC-ROIs (stage 2 and 3) in the HE-images by annotating all the cells that were positive for pHH3 and additionally annotated the unambiguous late phase MFs that failed to label for pHH3 (pHH3-assisted ground truth). In addition, a pHH3-assisted count was performed in the manually selected MC-ROIs (stage 1) if the MC of the respective participant was higher than in stage 2.

**Figure 2–3.**
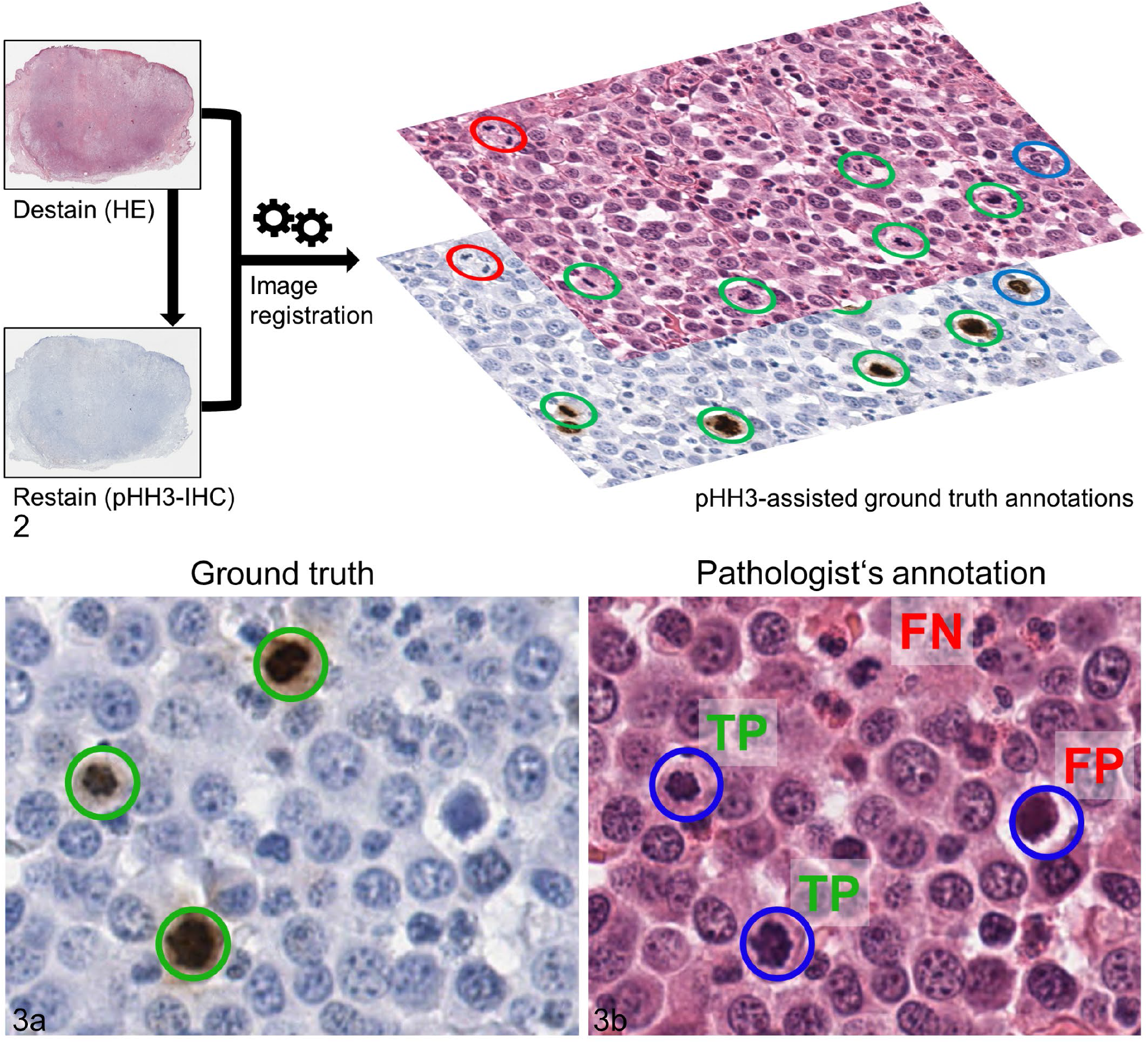
Immunohistochemistry-assisted ground truth. **Figure 2.** Labeling method of the ground truth dataset. The histological sections (hematoxylin and eosin stain, HE) were de-stained and re-labeled with immunohistochemistry against phosphohistone H3 (pHH3). Subsequently whole slide images of both staining methods were aligned on the cellular level via automated image registration and combined to decide if a tumor cell is a mitotic figure (MF) or not. Ground truth annotations comprised pHH3-positive cells that were recognizable on HE images (green circles) or were not readily identifiable on HE images (blue circles; especially prophase MF) as well as unambiguous late phase (especially telophase) MF that were pHH3-negative (red circles). Here these patterns are displayed as three distinct colors but in the ground truth dataset those structures were labeled as one label class. **Figure 3a.** High magnification image of an pHH3-stained tumor section with three positive tumor cells (green circles). **Figure 3b.** Histological image (HE stain) of the same tumor location than Fig. 2 with exemplary annotations by one of the study participants (blue circles). Compared to the pHH3-assisted ground truth, two annotations are true positives (TP), one is a false positive (FP) and one MF was missed (false negative, FN).

### Performance evaluation

The MC was defined as the number of annotations by participants within a MC-ROI. The pHH3-assisted MC was the number of ground truth annotations in the respective MC-ROI. Inter-observer agreement of the MCs between the three stages was calculated by the inter-observer correlation coefficient (ICC) and its 95% confidence interval (95% CI). ICC were evaluated as poor = 0–0.39, fair = 0.40–0.59, good = 0.6– 0.74, and excellent = 0.75–1.00.^11^ Differences were considered significant if the 95% CI did not overlap. The coefficient of variation (CV) between the pathologists at each stage was calculated.

Performance of classifying MCs into low and high according to the prognostic cut-off of MC ≥ 5 ^8,32,38^ was measured by accuracy as the number of correctly (below or above cut-off according to the pHH3-assisted ground truth) classified instances divided by all instances. To calculate the p-value for the difference in accuracy between Stage 2 and 3, a generalized linear mixed model (GLMM) was used. We fitted a logistic regression using correct classification (1 = correct, 0 = incorrect) as outcome and using a random effect for the pathologist to account for repeated measures (40 slides per pathologist).

For comparison of the manually selected approximate MC-ROIs, we produced images that visualize the region and a MC heatmap as an overlay on the WSI. The MC heatmap was based on the unverified algorithmic predictions and was calculated as previously described.^11^

The performance of the participants and algorithm to identify and classify individual MFs in the MC-ROIs of stage 2 and 3 (object detection task) was determined by standard object detection metrics.^10^ True positives (TP), false positives (FP) and false negatives (FN) were calculated against the pHH3-assisted ground truth (Fig. 3). A pathologist’s annotation and a ground truth annotation were counted as TP if both had a maximum Euclidean distance of 25 pixels (equivalent to 6.25 μm). True negatives are not available for object detection tasks.^10^ Precision (also known as positive predictive value), recall (also known as sensitivity) and the F1-score (harmonic mean of precision and recall) were defined as described by Bertram et al. ^10^: precision = TP/(TP+FP); Recall= TP/(TP+FN); F1 = 2× (precision × recall)/(precision+recall). Macro-averaged values were determined with the overall TP, FP, FN annotations/predictions of all cases totalled and thus every MF identification/classification did have the same weight regardless of the mitotic density of the case. This allows to evaluate the performance on the object level (each MF separately).^44^For the micro-averaged values, we determined the metrics (precision, recall, F1) for each slide and participant, and calculated the overall mean values at the end.^44^With this method, every case/slide has the same weight and the metrics represent the diagnostic performance on the sample level. 95% CI for the micro-averaged precision, recall and F1-score were determined using bootstrapping (5000 replicates, bias-corrected and accelerated CI). Differences between the object detection metrics were considered significant if the 95% CI did not overlap.

## Results

Twenty-tree of 26 (88%) pathologists from 11 laboratories completed this study. The remaining pathologists were excluded from analysis as they did not return results (N = 2) or did not examine all cases (N = 1). Of the included participants, 22 (96%) were Diplomates of the American (N = 12) or European (N = 10) College of Veterinary Pathologists and one (4%) had completed a residency in veterinary anatomic pathology. The duration of board certification ranged from 0 to 14 years (median of 6 years).

All 23 participants examined the 50 cases at the three examination stages (3,450 instances) and thereby created 38,491, 59,634 and 68,570 annotations in stage 1, 2 and 3, respectively. The number of annotations (all 50 cases combined) per participant for stage 1 ranged between 549 - 4,182 (mean: 1674; coefficient of variation, CV: 43%), for stage 2 between 1,263 - 4,412 (mean: 2593; standard deviation: 856, CV: 33%) and for stage 3 between 1,827 - 4,807 (mean: 2981; standard deviation: 638, CV: 21%). The pHH3-assisted ground truth dataset comprised 2,617 annotations (40 cases only) and the algorithmic predictions comprised 3,063 MF candidates (50 cases) for the ROIs of stage 2 and 3.

### Computer-assisted MCs have higher agreement

First, we evaluated the effect of computer assistance on the overall MC. The average MC for all cases and pathologists was 33.47 for stage 1, 51.86 for stage 2, 59.63 for stage 3 and 61.26 for the deep learning-based algorithm. Compared to stage 1, the average pathologist MC was increased by 54.9% in stage 2 and 78.2% in stage 3. Number of cases in which the MCs had very low values were notably reduced in stage 2 and 3. For example, MC = 0 were determined in 63 instances in stage 1 and only in 12 and 3 instances in stage 2 and 3, respectively (. Agreement of the MCs was higher between stage 2 and 3 (examination of the same MC-ROI) compared to stage 1 and 2 (mostly different MC-ROI). Inter-observer agreement of the MCs was good for stage 1 (ICC: 0.70; 95% CI: 0.60 - 0.79) and excellent for stage 2 (ICC: 0.81; 95% CI: 0.74 - 0.88) and stage 3 (ICC: 0.92; 95% CI: 0.88 - 0.96). The 95% CI of the ICC for stage 1 and stage 3 do not overlap; thus the difference was statistically significant. The CV for the MCs was 78.2% for stage 1, 51.3% for stage 2 and 34.9% for stage 3, thus reduced by more than half with full computer assistance. Figure 4 shows the improvement in agreement of the MCs with the pHH3-assisted ground truth MCs if computer assistance for identification/classification of MFs (stage 3 as opposed to stage 2) was available.

**Figure 4.**
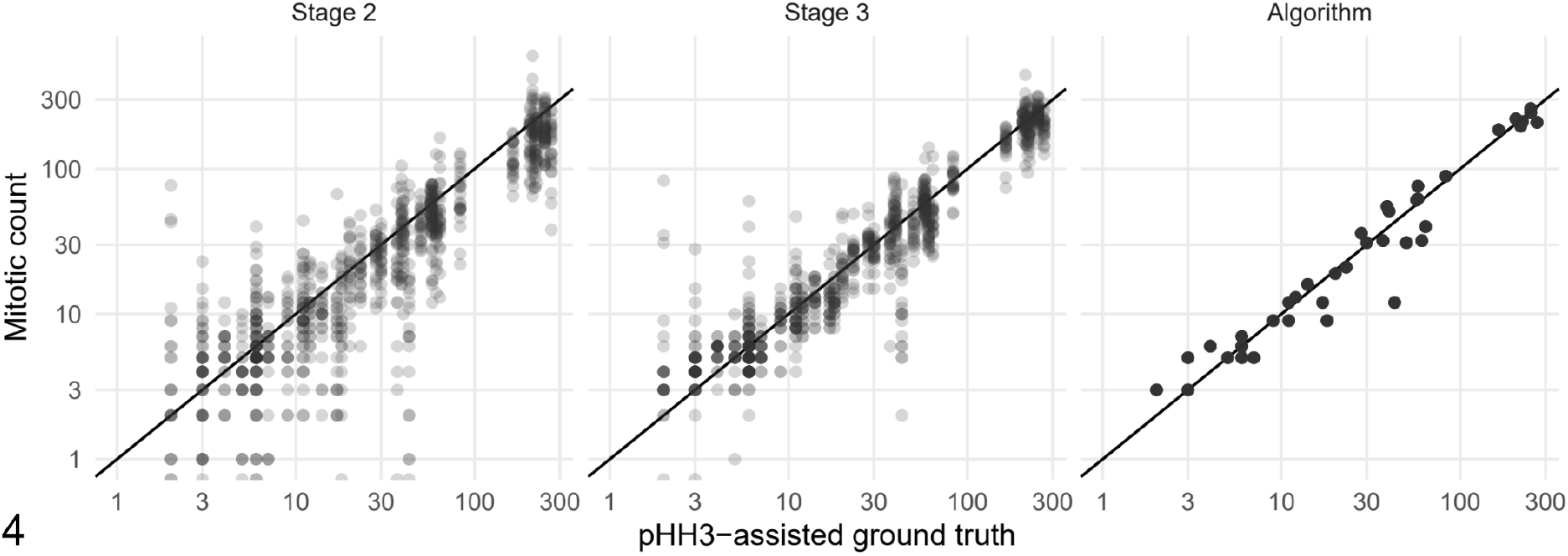
Scatterplots of the participant’s mitotic count (MC) values (stage 2 and 3) and the algorithmic (unverified) MC compared with the pHH3-assisted ground truth MC (all obtained in the same mitotic hotspot MC-ROI based on the algorithmic heatmap. The black line in the scatterplots indicate equal values for ground truth and pathologists or algorithmic MCs.

### Computer assistance improves accuracy of prognostic classification

Next, we evaluated how the computer-assisted approach influenced determination of values below and above the cut-off of MC ≥ 5 for tumor prognostication (published cut-off for the MC as a solitary prognostic parameter).^8,32,38^ For all pathologists and all 50 cases combined, 362, 170, and 116 instances were below and 788, 980, and 1034 instances above the cut-off in stage 1, 2 and 3, respectively. Compared to the pHH3-assisted ground truth MCs (available for 40 cases), accuracy of classification below or above the cut-off, respectively, was 75.8% (95% CI: 72.3% - 80.1%) for stage 1, 86.6% (95% CI: 84.2% - 89.7%) for stage 2, 91.7% (95% CI: 89.9% - 94.0%) for stage 3 and 95% for the deep learning-based algorithm. The increase of accuracy of the participants between stage 1 and 2 (p < 0.0001), stage 2 and 3 (p = 0.0012) and stage 1 and 3 (p < 0.0001) were significant. Due to the case selection strategy, the distribution of the hotspot MC values of the study cases is not representative of a routine diagnostic situation. We therefore divided the 40 cases into five tiers based on the pHH3-assisted ground truth MCs and determined individual accuracy values for these groups (Table 1). We determined that cases around the prognostic cut-off value had the lowest accuracy. However, also cases with a pHH3-assisted ground truth MCs between 25 and 49 had falsely low MCs in 20/138 instances (14.5%) without computer assistance (stage 1), while 9/138 (6.5%) and 1/138 (0.7%) instances of those cases were misclassified in stage 2 and 3, respectively.

**Table 1.** Accuracy of the 23 study participants and the deep learning-based algorithm to classify mitotic counts (MC) as below (MC < 5) or above (MC ≥ 5) the prognostic cut-off as compared to the pHH3-assisted ground truth MC (GT-MC).

| GT-MC | Number of cases | Accuracy for | Stage 1 * | Stage 2 | Stage 3 | Algorithm |
| --- | --- | --- | --- | --- | --- | --- |
| 0-4 | 4 | Below cut-off | 75.0% | 50.0% | 50.0% | 50% |
| 5-9 | 7 | Above cut-off | 31.7% | 70.2% | 82.6% | 100% |
| 10-24 | 8 | Above cut-off | 63.0% | 89.1% | 99.5% | 100% |
| 25-49 | 6 | Above cut-off | 85.5% | 93.5% | 99.3% | 100% |
| $\geq 50$ | 15 | Above cut-off | 99.4% | 100% | 100% | 100% |
| All cases | 40 | Below/above cut-off | 75.8% | 86.7% | 91.7% | 95% |

\* the GT-MC and the participants MC of stage 1 were not determined in the same tumor location. The GT-MC and the MC of stage 2 and 3 were determined in the mitotic hotspot location based on the algorithmic heatmap.

### Algorithmic area preselection is superior in finding mitotic hot spots

We evaluated the distribution of the manually selected MC-ROIs in stage 1 by the 23 participants. Visual assessment revealed that the approximate MC-ROIs were widely distributed throughout the tumor section in most cases, even if a high variability of the MC distribution (based on the algorithmic predictions) was present, i.e. mitotic hotspots were not detected consistently (Fig. 5–7). In only two cases was a similar tumor area was consistently chosen for the MC-ROI. In one case (No. 5; Fig. 8) a region within the tumor exhibiting local invasion was selected by all participants and in the other case (No. 37) the tumor location with highest cellular density was selected by 22/23 participants. This contrasts with automated image analysis that will always propose the same tumor area (100% intra-algorithmic reproducibility).

**Figure 5–8.**
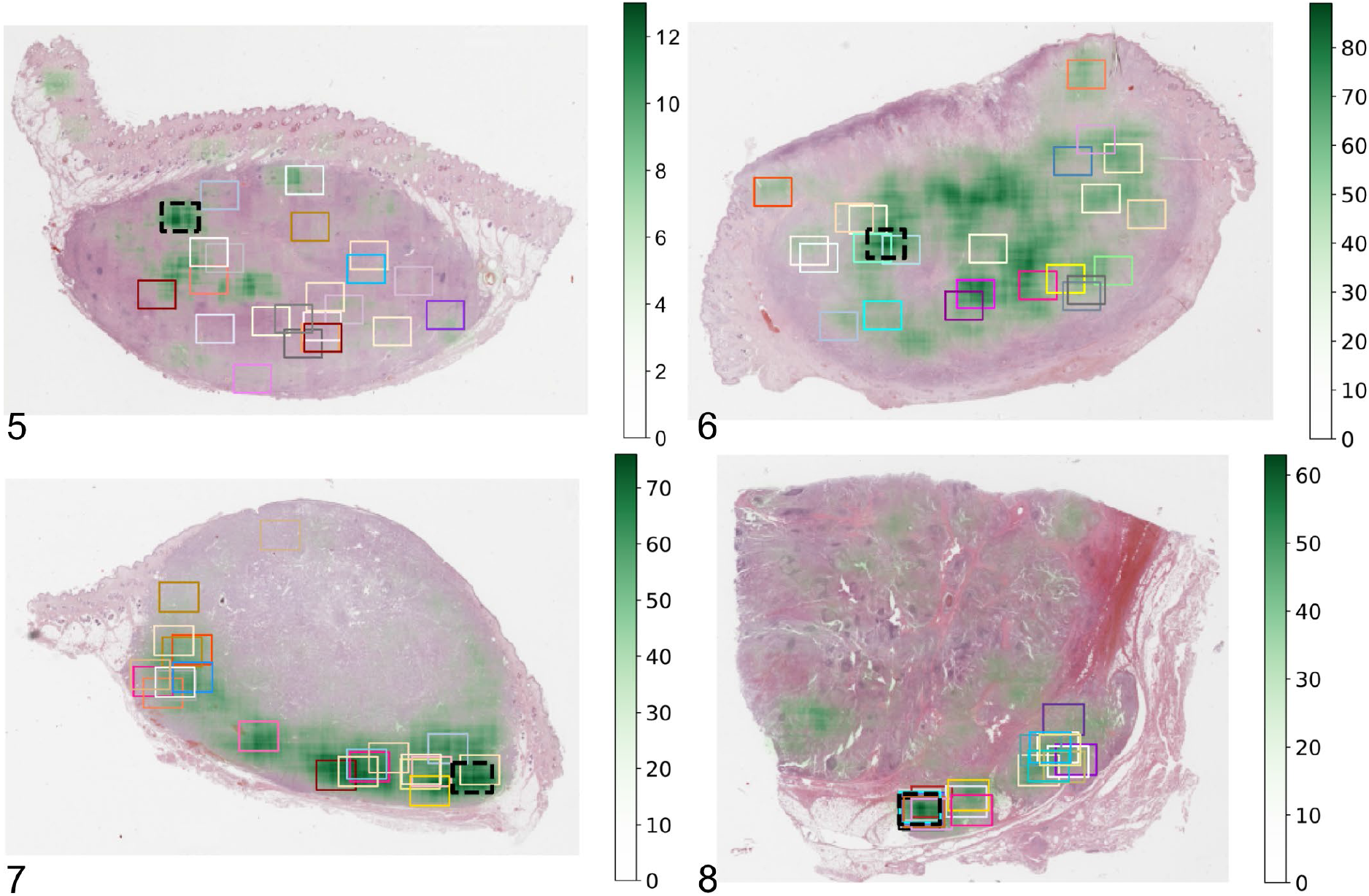
Approximate location of the mitotic count region of interest (MC-ROI) selected manually by each study participant (represented by the rectangular boxes) in the whole slide images. The black box with the dashed line represents the algorithmically preselected MC-ROIs. The estimated MC heatmap is visualized by variable opacity of a green overlay (scale on the right side of image) on the histological image (hematoxylin and eosin stain) and is based on algorithmic mitotic figure predictions. **Figure 5.** Case no. 33 with widely distributed MC-ROIs. Dark green areas represent mitotic hotspots. **Figure 6.** Case no. 46 with widely distributed MC-ROIs. **Figure 7.** Case no. 38 with MC-ROIs mostly along the tumor periphery. **Figure 8.** Case no. 5 with similar MC-ROIs at a side of local tumor invasion.

As it was the goal to find mitotic hotspots and we hypothesized that computer assistance is helpful for this, we compared the MCs of the participants in the manually selected MC-ROIs (stage 1) and algorithmically preselected MC-ROIs (stage 2). Of the 1,150 MC pairs from all participants, in 908 instances (79.0%) the MCs were higher in stage 2 and in 55 instances (4.8%) the MC was the same in stage 1 and 2. MCs of stage 1 were higher in 187/1,150 MC pairs (16.2%) for all participants combined and in 0 - 20/50 MC pairs (median: 8; mean: 8.1) for each individual participant. For 151/187 of those cases with higher MCs in stage 1, it was possible to perform pHH3-assisted MCs for both stages. Thereby we were able to prove that the mitotic density was truly higher in the manually selected MC-ROI in one instance (0.7%; ground truth MC = 4 and 3 in the two tumor locations), the same in both stages in 4 instances (2.6%) and actually higher in the algorithmically preselected MC-ROIs in 146 instances (96.7%). Only for case 27 the algorithmically preselected MC-ROIs was considered inappropriate as it comprised a tumor area with crush artifact that resulted in many false positive predictions (pHH3-assisted MC was not available for this case).

### Computer-assisted MF detection improves recall

The ability of the participants to identify and classify individual MFs was determined for the same MC-ROIs in stage 2 (unaided MF identification) and stage 3 (computer-assisted MF identification). Annotations of each participant and predictions of the algorithm were compared to the pHH3-assisted ground truth. For the participants, we found an overall decrease of FNs by 38.4% between stage 2 (23,107) and stage 3 (N = 14,256) and an overall increase of TPs by 23.7% between stage 2 (N = 37,117) and stage 3 (N = 45,929). An improvement of FNs and TPs was present for 22/23 participants and a negligible decline (by 3 annotations; <1%) was present for the participant that had the lowest number of FN and highest number of TN in stage 2. FPs only had an overall decrease by 3.8% between stage 2 (N = 10,981) and stage 3 (N = 10,582) whereas FPs were lower for 9 participants and higher for 14 participants. Subsequently, the macro-averaged recall had an overall increase by 14.6 percentage points (maximum 30 percentage points; 22/23 participants improved), the macro-averaged precision had an overall increase of 2.3 percentage points (maximum 21 percentage points; 11/23 participants improved) and the macro-averaged F1-score had an overall increase of 10.7 percentage points (maximum of 22.9 percentage points; 23/23 participants improved) between stage 2 and 3 (Table 2, Fig. 9–11). Differences between the two examination stages of the micro-averaged F1-score (12.4 percentage points), precision (5.2 percentage points) and recall (14.9 percentage points) were similar (Table 2). 95% confidence intervals for the micro-averaged F1-score, recall and precision did not overlap between stage 2 and 3 (Fig. 9); thus the improvement is significant. The deep learning-based algorithm (unverified predictions) had an almost balanced proportion of FP detections (N = 406) and FN detections (N = 496) and subsequently similar values for precision (0.84), recall (0.81) and the F1-score (0.83). The overall performance (F1-score) of the algorithms was comparatively high as it was not reached by participants in stage 2 and was only slightly exceeded by two participants in stage 3 (Fig 11). The F1-score of the algorithm was significantly better than the score of the participants of both stages (CI did not overlap).

**Table 2.** Performance (macro- and micro-averaged metrics with range or 95% confidence interval, CI) of the 23 participants (partially or fully computer-assisted of stage 2 and 3, respectively) and the deep-learning-based algorithm (unverified predictions) for detecting individual mitotic figures in in mitotic hotspot regions of interest compared to a pHH3-assisted ground truth.

| Metrics | Precision |  | Recall |  | F1-score |  |
| --- | --- | --- | --- | --- | --- | --- |
|  | 2 | 3 | 2 | 3 | 2 | 3 |
| Examination stage | 2 | 3 | 2 | 3 | 2 | 3 |
| Participants, macro-averaged values (range) | 0.80 (0.56 - 0.95) | 0.83 (0.58 - 0.94) | 0.62 (0.37 - 0.82) | 0.76 (0.54 - 0.87) | 0.68 (0.53 - 0.79) | 0.79 (0.67 - 0.84) |
| Participants, micro-averaged values (95% CI) | 0.74 (0.72 - 0.76) | 0.79 (0.78 - 0.81) | 0.62 (0.60 - 0.63) | 0.76 (0.75 - 0.78) | 0.63 (0.61 - 0.64) | 0.75 (0.74 - 0.77) |
| Deep learning-based algorithm, macro-averaged value | 0.84 |  | 0.81 |  | 0.83 |  |
| Deep learning-based algorithm, micro-averaged value (95% CI) | 0.83 (0.80 - 0.85) |  | 0.80 (0.76 - 0.84) |  | 0.80 (0.77 - 0.82) |  |

**Figure 9–11.**
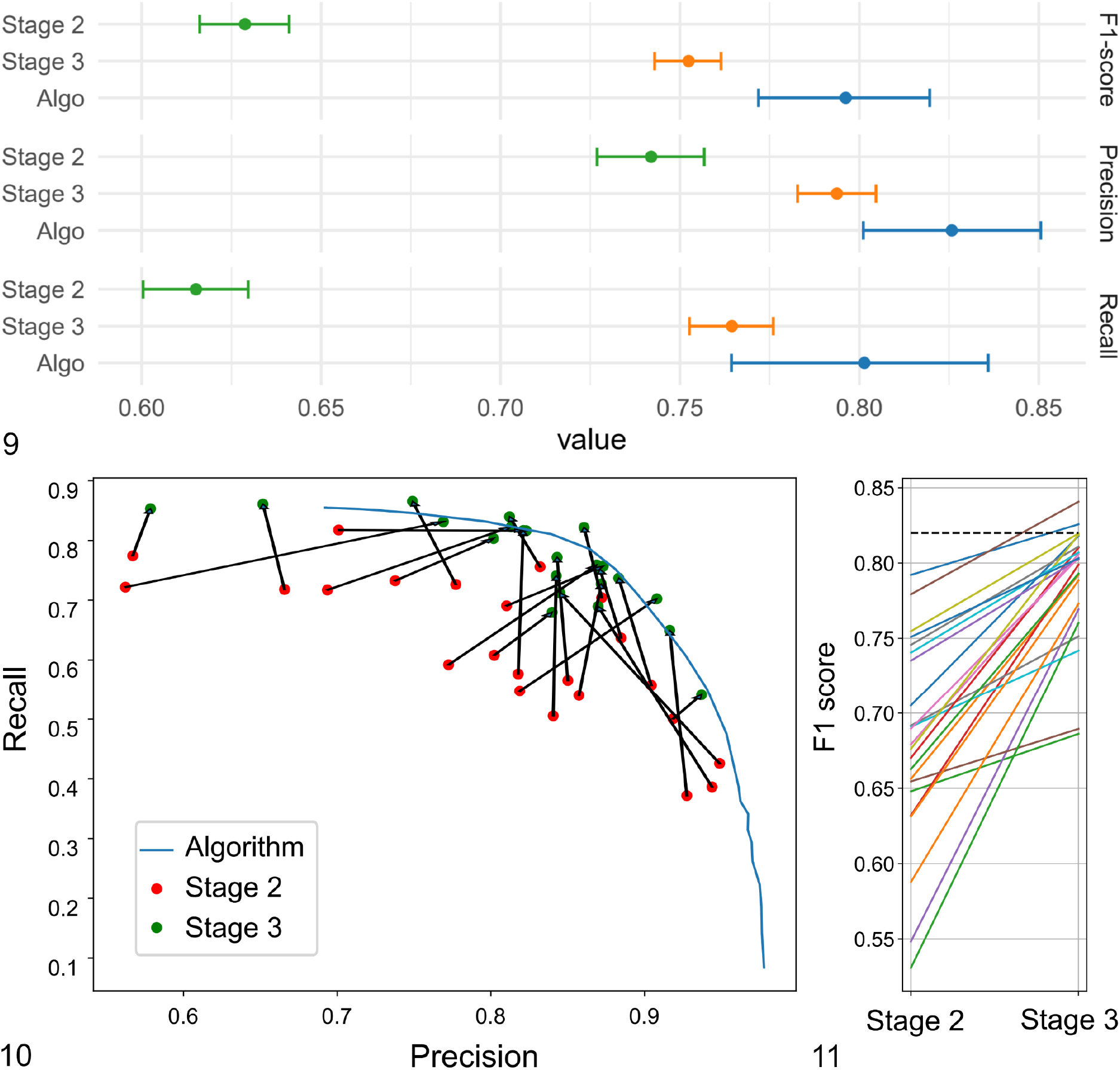
Object detection performance (identification and classification of mitotic figures) of the 23 participants and the deep learning-based algorithm in stage 2 and 3. **Figure 9.** micro-averaged F1-score (upper graph), precision (middle graph) and recall (lower graph) with their 95% confidence intervals. The difference is considered significant (p<0.05) if the intervals do not overlap. **Figure 10.** Recall and precision for the individual participants in stage 2 and 3 (connected by a black arrow) and the precision-recall-curve for the algorithm (at different classification thresholds. For the algorithmic predictions for the present study, a single classification threshold was used that resulted in a recall of 0.81 and precision of 0.84. **Figure 11.** F1-scores for the individual participants for stage 2 and 3. The dashed black line represents the F1-score of the algorithmic predictions.

### Participants considered algorithmic MC-ROI preselection to be the most helpful feature

Twenty-one of the 23 study participants (91%) filled out the concluding survey after they finished stage 3. Analysis revealed that participants considered MC-ROI selection to be the most difficult aspect of performing the manual MC (stage 1) whereas spotting potential MF candidates and classifying them against look-alikes was considered comparably easy. Subsequently most participants had the subjective impression that algorithmic MC-ROI preselection was extremely or very helpful and visualization of algorithmic predictions as well as display of their algorithmic confidence values was generally considered helpful to a lesser degree. Almost all participants (N = 20 / 21) indicated that their decision of classifying MFs against look-alikes was consciously influenced in stage 3 by the algorithmic confidence value at variable degrees ranging from being influenced in very few to many potential MF candidates, especially if participants were uncertain about the MF candidate (N = 17 / 20). Most participants (N = 15 / 21) considered digital microscopy generally inferior (to a variable degree) to light microscopy for identification of MF and mostly deemed fine-focusing / z-stacking necessary for at least some MF candidates (N = 14 / 21). The majority of the participants (81%; N = 17 / 21) considered the rectangular shape of the MC-ROI to be acceptable for performing the MC in the examined cases, while two participants found the shape inappropriate (two participants had no opinion).

## Discussion

The present study confirmed that inconsistency and inaccuracy of the MC arises from a combination of inappropriate MC-ROI selection (failure to find mitotic hot spots), incomplete MF identification and imprecise MF classification. We addressed all three aspects with our computer-assisted MC approach via algorithmic area preselection (stage 2), MF candidate visualization (stage 3) and display of algorithmic confidence values (stage 3). We were able to prove significantly increased performance on the overall MC-level and individual MF-level with computer assistance. Participants of the present study reported that screening tumor sections for mitotic hotspots is the most difficult task of manual MCs and subsequently deemed algorithmic MC-ROI preselection by far the most useful tool. In fact, we have shown that algorithmic area preselection was superior to manual area selection in almost all instances. Nevertheless, visualization of potential MF candidates also had a strong positive effect on the pathologists’ ability to find MFs in MC-ROIs (recall). In contrast, the benefit of displaying the algorithmic confidence values was considered controversial by participants (see below) and did not seem to have a strong positive impact on precision. It was beyond the scope of the present study to determine the prognostic value of computer-assisted MCs, which should be done in future studies. While we have shown a more consistent classification of the cases into prognostic cut-off values (in our case MC < 5 and ≥ 5 ^8,32,38^) with computer assistance, we have also found an overall higher MC for our computer-assisted methods. Therefore, new prognostic cut-off values and stratifications have to be determined (likely for each individual computer-assisted approach / software tool). For example, Elston et al. ^18^ have proposed a cut-off of 0 for group 1 ccMCTs with good prognosis (group 2: 1-7; group 3: > 7). This prognostic group is notably reduced in larger tumor sections (by a factor of 21x) if full computer assistance (stage 3) is used due to the improved sensitivity of MF detection. A limitation of the present study is that only a few cases with a truly low MC (below cut-off value) were included and future studies need to verify our results for this subgroup.

A particular strength of the present study was the use of a pHH3-assisted ground truth for performance evaluation. Most previous studies have evaluated their MF algorithms against a majority vote of pathologists annotations from HE images,^3,5,12,30,35,41,43^ which might be problematic due to human limitations in performing this task. If a consensus is generated by such a large group of participants as in the present study, a majority vote is very likely to include predominantly clear MFs and would not contain many morphologically inconclusive or equivocal, but still “true” MFs. Hence, a majority vote will not necessarily represent the biological truth.^10^ The advantage of pHH3 immunohistochemistry (as opposed to HE images) is that MFs are easily spotted (high sensitivity) and much more easily classified against look-alikes (high specificity with the exception of telophase MF).^16,19,33^ We highlight that some of the annotated pHH3-positive cells were extremely difficult to classify as MFs in the HE image (especially early prophase or if cells were tangentially sectioned) and might therefore have been missed (false negative) by study pathologists in all MC approaches. It is acknowledged that pHH3-derived labeling may overestimate prophase MFs and underestimate telophase MFs as compared to solely HE-based labeling.^36^ Therefore, we decided to use a combination of the pHH3 and HE image for annotating the ground truth. We believe that our ground truth labeling approach is the most objective and accurate reference method available for MFs to date and was independent of the participants (as opposed to a majority vote ground truth).

We have shown that identification of mitotic hotspots can be tremendously improved with computer assistance.^5^ An advantage of algorithms is that they can efficiently analyze entire WSIs with 100% intra-algorithmic reproducibility and are therefore able to determine the mitotic distribution consistently in entire or even multiple tumor sections. A limitation of our deep learning method was that 10% of the cases analyzed had an algorithmic MC-ROI preselection outside of the tumor area and one case had an algorithmic MC-ROI preselection in an area of crush artifacts. A previous study reported inappropriate MC-ROI selection in 58% of the cases.^7^ While it was beyond the scope of the present study to investigate approaches to resolve this problem, we propose that computer-assisted MC tools include one of the following features: 1) restriction of the algorithmic predictions to the actual tumor area, which can be delineated manually by a pathologist ^7^ or segmented automatically by means of deep learning-based algorithms;^20^ 2) heat map visualization of mitotic density for manual selection / correction of the MC-ROI; 3) proposal of the top 3-5 hotspots areas from which pathologists can choose. Another aspect that needs to be considered for future implementation of computer-assisted approaches is the shape of the MC-ROI. Even though most participants found the rectangular shape appropriate for the examined ccMCT cases, it might possibly be useful to be able to adjust the MC-ROI shape for tumor types that have a highly variable cellular density and/or may include a variable high proportion of non-neoplastic tissue, that should be excluded from the MC-ROI.^17,29^

Even if pathologists examine the same image sections, variability of the number of enumerated MFs has been noted ^15,40,41,43^ and computer assistance seems to be a promising solution for improving MF identification/classification. Similar to the comparison between stage 2 and 3 of the present study, Pantanowitz et al. ^30^ investigated the influence of computer assistance on the pathologist’s performance of annotating MFs in “hpf” images. Pathologists in this study ^30^ had a somewhat lower performance (F1-score) overall as compared to our results (7.1% lower without and 7.7% lower with computer assistance), which might be related to the ground truth definition used. Pantanowitz et al. ^30^ used the majority vote of 4/7 pathologists to define a “true” MF (see above). Nevertheless, Pantanowitz et al. ^30^ demonstrated a similar increase of the overall F1-score as in the present study (10% vs. 10.7%); however, they had a slightly lower increase of the overall recall (11.7% vs. 14.6%) and higher increase of overall precision (8.8% vs. 2.3%) than our study. This might also be related to the ground truth definition used, or the performance of the image analysis algorithm applied, or due to the level of the individual pathologists’ acceptance of the algorithmic predictions. Of note, our results demonstrate that individual pathologists may have high recall / low precision, moderate recall / moderate precision, or high precision / low recall. This is probably influenced to a large part by the decision criteria of individual pathologists for ambiguous patterns, and standardized morphological criteria for MFs might be helpful to harmonize the pathologist’s decision.^17^ Although all pathologists had overall a high performance (F1-score), the individual precision-recall-tradeoff of different pathologists may have a tremendous influence on the MCs and the prognostic stratification based on specific cut-off values. Our results show that the computer-assisted approach (stage 3) generally shifted the individual pathologists somewhat towards a more harmonic trade-off between recall and precision (moderate recall/moderate precision). The overall direction of the shift coincides with the precision/recall of the algorithmic results and could be influenced by a confirmation bias of the experts. Unlike the study by Pantanowitz et al. ^30^ we supplied the study participants with the algorithmic confidence value in stage 3. Although the model confidence scores overall had a strong correlation with the ground truth, some participants commented that a few confidence values contrasted their degree of certainty. Interestingly, many participants felt that they were (negatively) biased by the confidence values especially for difficult MF candidates. A previous study has in fact shown that pathologist may fail to identify a high proportion of incorrect (false negative and false positive) algorithmic predictions ^26^ and future studies need to determine the (positive or negative) effect of this bias in a diagnostic setting.

The results of our study were highly dependent upon adequate performance of the applied algorithm. Algorithms always exhibit 100% intra-algorithmic reproducibility, but accuracy may vary largely between different algorithms (inter-algorithmic variability; depending on numerous factors). The algorithm used in this study was created and evaluated on a distinct dataset without employing IHC labeling. Yet, the algorithmic predictions of the present study were highly accurate (F1-score) compared to the pHH3-assisted ground truth (as compared to the study participants). The high performance of the algorithm used for the present study can be attributed to the advanced deep learning methods applied,^5^ the high quality and quantity of the dataset used for training of the algorithm ^12^ and the high representativeness of the training dataset for the present study cases. However, we also experienced several incorrect MF predictions and few inappropriate algorithmic MC-ROI preselections, which necessitates review by trained experts (computer-assisted MC) and further research on algorithmic solutions.

One of the most relevant concerns that cannot be considered solved to date is robustness in the application of such algorithms on images that are acquired with other scanners or have strong differences in HE staining (domain shift).^3,4,10^ The consequence is that algorithms cannot necessarily be applied to images from different laboratories without prior validation and possibly modification (such as by color normalization and augmentation ^36,40^, domain adaptation ^4^ and/or transfer learning ^3^). A limitation of deep learning-based models is that the specific decision criteria are mostly not transparent (“black box”) and situations in which the algorithm may fail are not easily foreseeable. It is those “unexpected” situations that highlight the necessity of careful validation of each new application for its intended use. Deep learning models can be tested statistically (using metrics such as recall, precision or the F_1_ score),^10^ and each laboratory that wants to use a MF algorithm should validate its performance based on those metrics. The 50 cases included in the present study support the impression that the algorithm works reliably in many cases, yet the included cases may not represent all potential sources of error. Therefore, we consider computer-assisted approaches with verification by a trained pathologist to be highly beneficial; at least until more experience and progress is gained through research and routine use of these applications. Access to intermediate results of the algorithmic pipeline (such as visualization of MF predictions) may allow a higher degree of comprehensibility by the reviewing pathologist and should therefore be encouraged. Future studies need to determine which degree of expert review is required, how reliably pathologists can detect algorithmic errors and to what degree the decision of pathologists is consciously and unconsciously biased by algorithmic predictions.

## Conclusion

Our results demonstrate that computer assistance using an accurate deep-learning based model is a promising method for improving reproducibility and accuracy of MCs in histological tumor sections. Full computer assistance (assistance in MC-ROI selection and MF identification/classification) was superior to partial computer assistance (only assistance in MC-ROI selection) in the present study. This study shows that computer-assisted MCs may be a valuable method for standardization in future research studies and routine diagnostic tumor assessment using digital microscopy. Furthermore, improved work efficiency (such as by MC-ROI preselection) may be of interest for diagnostic laboratories and additional studies need to evaluate the degree of verification of algorithmic predictions required. Future research should also evaluate whether computer-assisted MC approaches will benefit tumor prognostication (compared to patient outcome).

## Acknowledgement

We thank Nicole Huth for technical support.

